# Structure of the 5’ untranslated region in SARS-CoV-2 genome and its specific recognition by innate immune system via the human oligoadenylate synthase 1^†^

**DOI:** 10.1101/2021.12.10.472066

**Authors:** Emmanuelle Bignon, Tom Miclot, Alessio Terenzi, Giampaolo Barone, Antonio Monari

## Abstract

2’-5’-Oligoadenylate synthetase 1 (OAS1) is one of the key enzymes driving the innate immune system response to SARS-CoV-2 infection whose activity has been related to COVID-19 severity. In particular, OAS1 is a sensor of endogenous RNA that triggers the 2’-5’ oligoadenylate/RNase L pathway in response to viral infections, ultimately activating the RNA-Lyase which cleaves endogenous and exogenous RNA impeding the viral maturation. Upon SARS-CoV-2 infection, OAS1 is responsible for the recognition of viral RNA and has been shown to possess a particularly high sensitivity for the 5’-untranslated (5’-UTR) RNA region, which is organized in a double-strand stem loop motif (SLA). Since the structure of the nucleic acid/protein complex has not been resolved, here we report its structure obtained by molecular modeling, including enhanced sampling approaches. We also pinpoint that the SL1 region enhances the interaction network with the enzyme, promoting specific hydrogen bonds, absent in normal double strand RNA fragments, hence rationalizing the high affinity for OAS1.

---

Upon sensing of endogenous double-stranded RNA, OAS1 catalyzes the formation of the secondary messenger 2’-5’ oligoadenylate, which subsequently activates the RNase L enzyme responsible for RNA cleavage, thus stopping viral replication. As a matter of fact, OAS1 plays an important role in the response to SARS-CoV-2 infection. Indeed, the severity and outcome of COVID-19 have been linked to OAS1 polymorphisms^1–3^, making it an inter-esting target for antiviral drugs development^4,5^. OAS1 is sensitive to RNA sequence^6^, and has been proposed to have a strong affinity for the 54 first nucleotides of the 5’ untranslated region (5’-UTR) of SARS-CoV-2^2^. The characterization of the secondary structure of this RNA region indicates that it is organized in two distinct stem loops, of whom the first one (SL1) exhibits a remarkably high affinity for OAS1^7,8^. Furthermore, the SL1 structure a ? a crucial role in regulating the genome replication, and particularly the action of the RNA dependent RNA polymerase.^9,10^ Models of the SL1 structure have been reported, based on secondary structure predictions^11,12^, and more recently on experimental data^13^. However, there is no tertiary structure of the SL1 ds-RNA interacting with OAS1, which would provide crucial atomic-scale information.^14^ Here, we describe the tertiary structure of the OAS1/SL1 complex as well as its dynamical behavior resolved using protein/nucleic acid docking, all-atom equilibrium molecular dynamics (MD) simulations and its Gaussian Accelerated extension (GAMD). The characterization of the interaction network between SL1 and OAS1 highlights the importance of the hairpin organization in promoting the high affinity of the immune system protein for this specific RNA fragment. Hence, our work may provide important molecular basis for antiviral drug development, specifically acting against SARS-CoV-2 infection and targeting the OAS1-RNAse L pathway. The possibility of targeting the UTR genome regions is also attractive due to the fact that this region appears fundamental to finely regulate RNA replication.^15^

However, before analyzing its interaction with OAS1 the native structure of SL1 needs to be resolved. To this aim we firstly generated the an initial structure of SL1 based on its sequence using Unafold web server.^16^ Our 3D model, spanning the first 40 residues of the 5’-UTR sequence, shows the spatial organization of the rG7-rC33 in a SL motif presenting a central bulge due to unpaired residues rA12 and rA27-rC28. The double-stranded region is connected by a loop (residues rU18-rC21) at its extremity – see Figure 1.

**Fig. 1.**
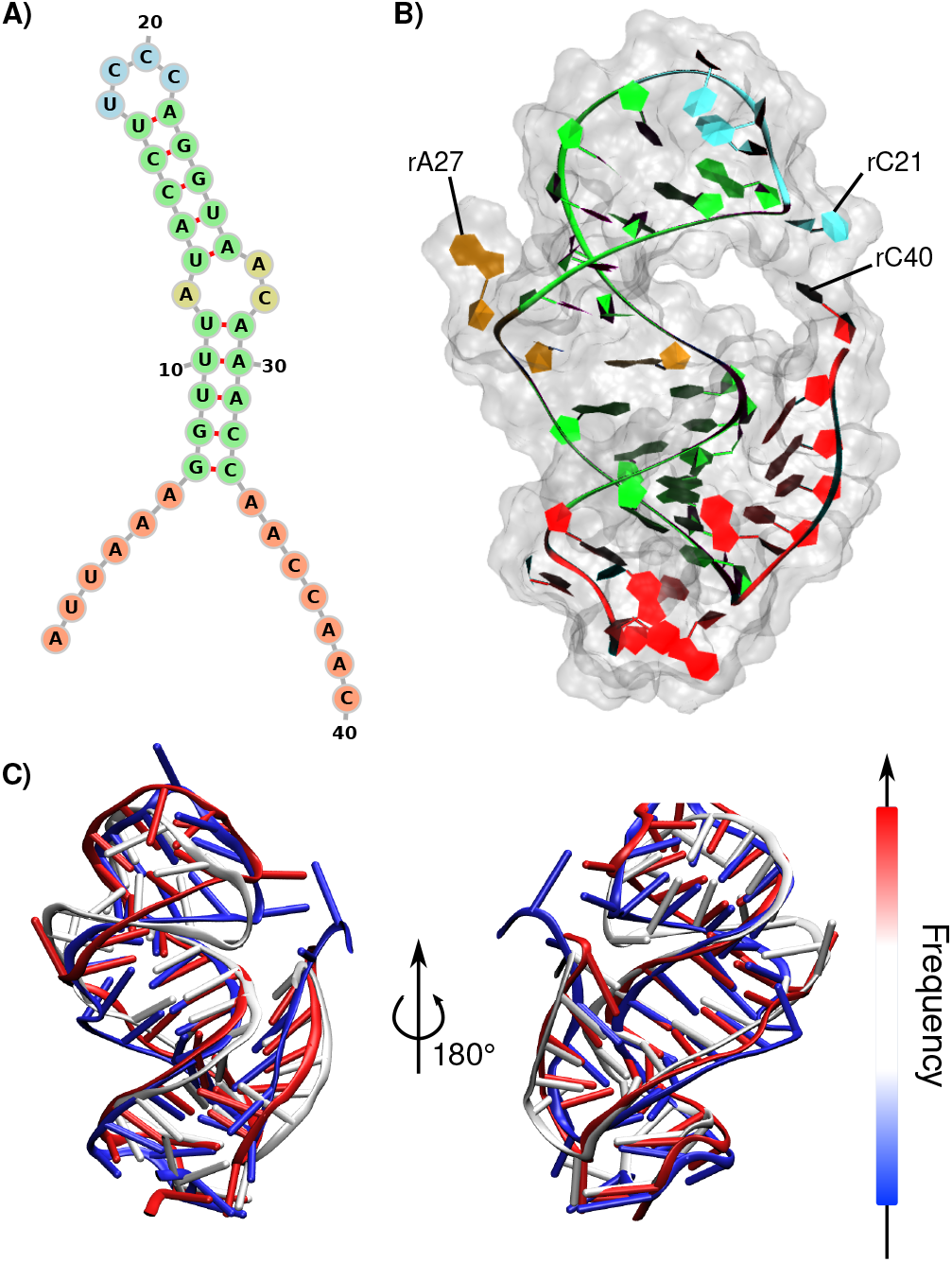
A) Secondary structure of the 5’-UTR SL1 region of SARS-CoV-2 genome and B) a representative structure of the reconstructed 3D model. The loop, bulge, double-stranded, and single-stranded regions appear in blue, brown, green and red, respectively. C) Superimposed representative structures of the three major clusters extracted from the MD trajectory of the SL1 region.

To assess the stability of the structure, unrestrained *μ*s-scale MD simulations have been performed revealing the monotonous conformational behavior of the SL1 structure. Indeed, the loop and to a lesser extent the bulge appears the only flexible regions. Our results are globally coherent with those obtained by Bottaro *et al.*^13^ However, differently from the precedent studies, we found that, while rA27 is most frequently excluded from the helix, rC28 remains,in the major groove 60% of the simulation time, due to interactions with the facing rA12. Yet we also observe transient conformations exhibiting an extruded rC28 and a helical rA27. These differences might be due to the presence of the 5’ and 3’ single-strand extremities in our model, which interestingly fold onto the SL structure and interact with its backbone.

Additionally, the transient folding of the upper region of the SL onto the double-stranded region, which is observed in ca. 25% of the trajectory, allows favorable interactions between rC21 and the rA12 backbone, hence assisting the extrusion of rC28. The rest of the time, the extruded rC21 is instead stabilized by *π*-stacking with 3’-rA39 and rC40. The presence of extensive hydrogen bond network, involving the loop, the bulge, and the 5’ single stranded regions imposes a strong bend of 104.3 ± 34.6 ° to the SL structure, as shown by Curves+^17^ analysis – see Table 1.

**Table 1.** Total bend angle of the RNA for each system.

| system | average (°) | stdev (°) |
| --- | --- | --- |
| SL1 | 104.3 | 34.6 |
| OAS1-SL1 | 113.0 | 28.5 |
| OAS1-RNA crystal | 18.8 | 8.7 |

The Root Mean Square Deviation (RMSD)-based cluster analysis of the MD ensemble confirms that the loop is the most flexible region of SL1, while the rest of the sequence exhibits a rather stable structure – see Figure 1-C. As expected, the single-stranded regions also exhibit a pronounced flexibility, despite developing important interactions with the double-stranded region. while being interesting *per se,* the characterization of the SL1 conformational space provides crucial information, notably a starting structure of the RNA fragment, for the study of its complexation with OAS1, as described in the following.

The most populated SL1 conformation, extracted from the MD ensemble was docked, using the HDock web server,^18^ on the crystal structure of OAS1, which has been retrieved from PDB 4IG8^19^. The resulting structure suggests that the 5’ minor groove of SL1 develops contacts with the OAS1 N-lobe while the second RNA minor groove, together with the loop region, are oriented towards and interact with the protein C lobe. On top of this initial structure we have performed equilibrium MD simulations using the NAMD code. ^20,21^ As detailed in the Supplementary Information the RNA-specific *χ_OL3_* force field^22^ has been used together with the AMBER FF14SB for the description of the protein.

Molecular dynamics simulations of the OAS1/SL1 complex, and of the reference crystal structure complexed with ideal double-stranded RNA,^19^ highlights different interaction network in the two systems. Indeed, the strong bending of SL1, i.e. 113.0 ± 28.5 °, is even more pronounced upon complexation with OAS1 than for the isolated strand, see Table 1. This and the presence of the SL induce a very specific binding mode.

The two accessible minor groove regions of SL1 anchor the RNA to the OAS1 surface (see Figure 2), with hydrogen bonding between the RNA backbone/sugar and K42, R195, K199, and T203 of OAS1, similarly to what is observed for ds-RNA (Figures S1 and S2). However, the interaction network is more extended in the reference ds-RNA/OAS1 than for SL1. The hydrogen bonds, evidenced for the crystal structure,^19^ are persistent throughout the simulation, highlighting the non-specific interactions with ds-RNA backbone and sugar moieties (Figure S3). This might be due to the more rigid and less curve structure of the reference ds-helix, which exhibits an average bend of only 18.8 ± 8.7^°^.

**Fig. 2.**
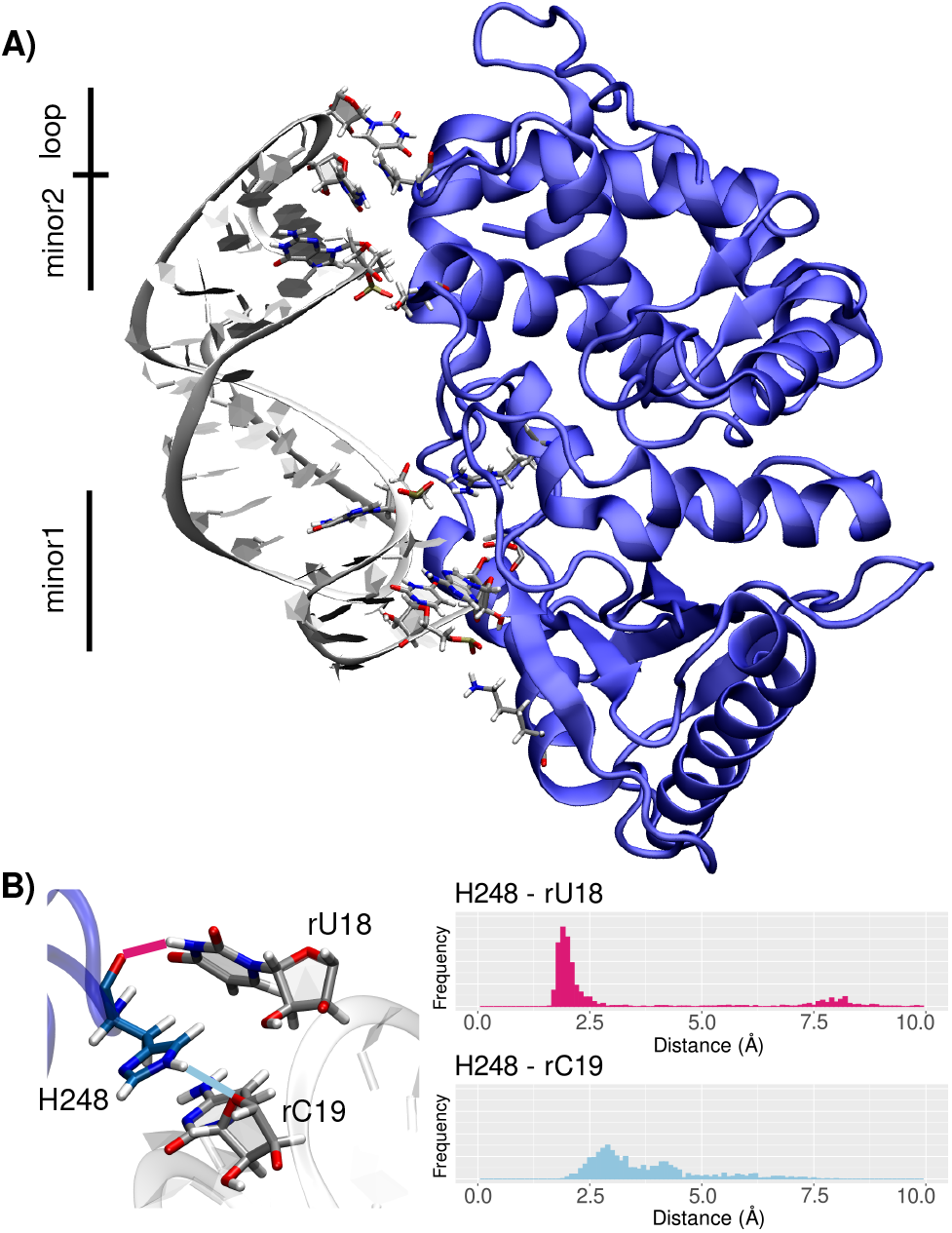
A) Representative structure of the OAS1/SL1 complex exhibiting contact surfaces around the two minor groove and the RNA hairpin (loop). SL1 appears in white and OAS1 in blue, and the interacting amino and nucleic acids are displayed in licorice. B) Key-interactions in the loop region between H248:O and rU18:H3 in magenta, and between H248:HE2 and rC19:O4’ in light blue (left), and their distribution along the MD simulation (right).

Our MD simulations allow to highlight the sequence dependency of the SL1 recognition by OAS1. Indeed, its specificity relies on the interactions with the RNA nucleobases. In both the initial crystal structure and the simulations of the reference ds-RNA system, specific hydrogen bonds involve S56 backbone, Q158, K42, and T203. On the other hand, the specific interactions of SL1 through the nucleobases involve different residues, which are mainly spanning the extruded RNA bases and the loop region:

T247 interacts with rC19, and V58 and H248 backbone atoms interact with rA6 and rU18. The interaction of H248 with the RNA loop appears particularly important for SL1 binding to OAS1. Indeed, H248 is involved in a very persistent hydrogen bonds with rU18 through its backbone and with rC19 through its side chain (Figure 2). Consequently, the OAS1/SL1 complex exhibits three important contact surfaces involving the loop and minor grooves of SL1, and both N and C lobes of OAS1. Interestingly, the bulge region of SL1 is not involved in the interaction network with the protein. The shift from rather unspecific backbone-driven to nucleobase-centered interaction may explain the affinity of OAS1 for the SL1 sequence, the question of the conservation of its sequence under the evolutionary pressure is till to be addressed. Miao *et al.* ^7^ have pointed out that the extruded bases on the bulge can be involved in base pairs in some variants, however their limited participation to the recognition should not hamper selectivity. As concerns the loop region, while the 5’-UTR and the stem-loop arrangement appear as fundamental for the viral replication^15^, the SL1 region has nonetheless been recognized as a hotspot for point mutations. However the most common allele modifications involve the quartet rA34, rA35, rC36, and rC37^15^, which seems less involved in OAS1 recognition.

To ensure that our MD simulations provided a complete exploration of the conformational space of the OAS1/SL1 complex, and most notably describe its most stable conformations, a Gaussian-Accelerated MD (GAMD) run,^23,24^ in which an energetic repulsive bias is applied to all the dihedral angles of the protein and the nucleic acid, was performed to enhance the sampling of the phase space and avoid local minima traps. To take care of the effect of the bias, the re-weighted free energy map was obtained as a function of the projection of the MD trajectory on top of the two main RNA Principal Component Analysis (PCA) vectors, used as collective variables (Figure 3). The two main PCA vectors largely dominate the expansion, and as seen in Figure S5 while the first vector (PCA1) mainly describes the collective detachment of RNA from OAS1, the second one (PCA2) involves the bending and compression of SL1. From the analysis of the re-weighted free energy map one can evidence three main minimum basins. Interestingly, the principal basin represents a rather extended and flat energy surface which globally spans values of PCA1 comprised in the −40/ + 40 interval, showing only moderate energy penalties not exceeding 10 kcal/mol. The other two basins are, on the other hand, separated by higher barriers and are globally much less extended than the principal one. By analyzing the representative structures belonging to the different basins, one can see that the main features already evidenced by equilibrium MD simulations are indeed preserved. In particular the main contact regions and interaction network, are globally maintained, involving the minor groove region and the extruded bases. On the same level the strong curvature of the RNA fragment is also maintained as well as the role of the free nucleobases interaction with the enzyme. Indeed, the persistence of the H248 interaction with the RNA bases located in the SL loop should be underlined. Likewise, the loop region experiences some flexibility, and constitutes the area more subjected to structural variation. This backbone deformations are driven by the tendency to maximize hydrogen-bonds and polar interactions between the dangling nucleobases and the polar or charged OAS1 residues as shown in Figure 3.

**Fig. 3.**
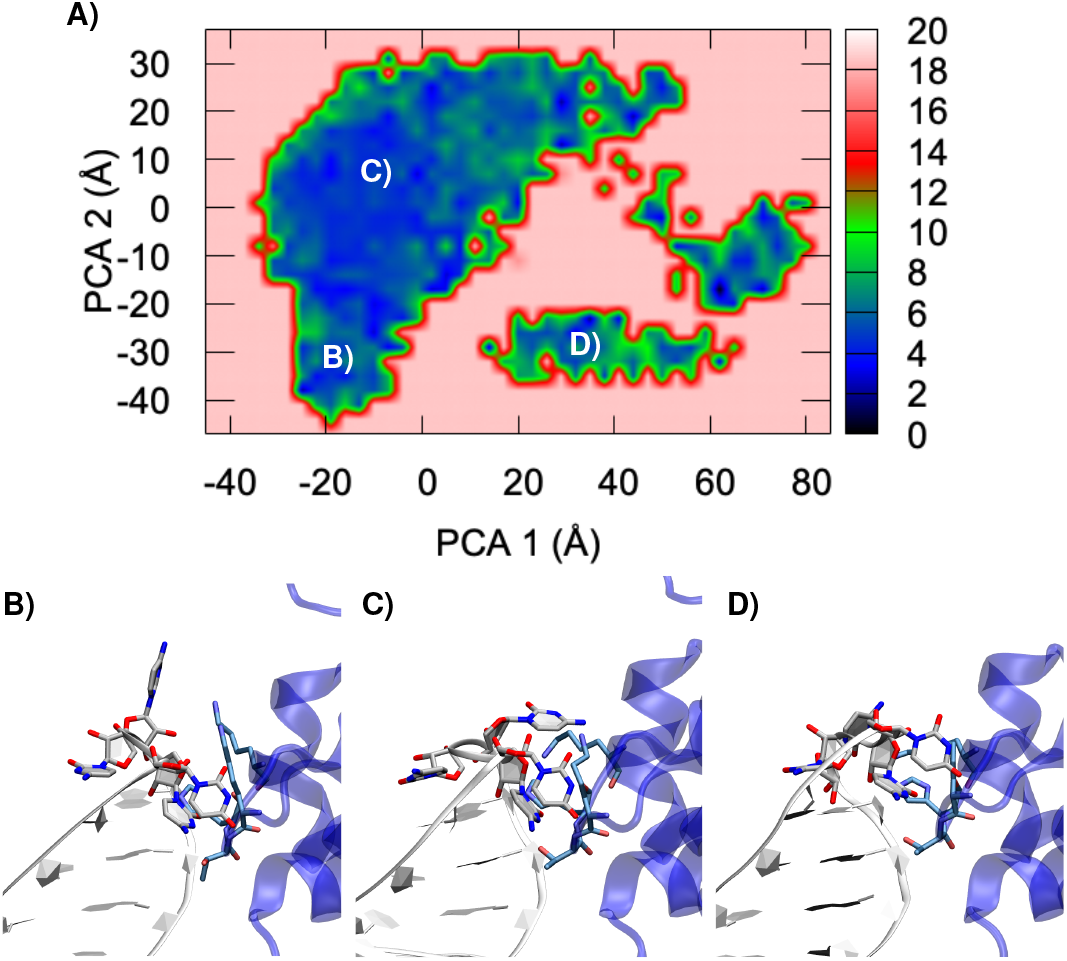
A) Re-weighted free energy map obtained from the GAMD analysis as a function of the two main PCA modes. (B,C, and D) representative snapshots of the minimum regions, in which we have highlighted the loop area and the interacting residues, their location in the free energy map is also highlighted (see global structures in Figure S5).

The response of the immune system to infections relies on the precise recognition of exogenous genetic or proteic material. The complex regulation of the innate immune system response to infections has been brought on the front line by the outbreak of the COVID-19 pandemics. Indeed, while an efficient immune response is needed to counteract the effects of SARS-CoV-2 infection, its deregulation has also been recognized as the cause of serious outcomes and morbidity. The recognition of exogenous viral RNA by the OAS1/RNAse L pathway is of particular importance in the innate response to SARS-CoV-2 infection. ^25^ Indeed, it has been shown that variants of OAS1 inducing colocalization of the protein in the viral replication regions correlate with milder symptoms and better systemic response. Furthermore, a high *in vitro* affinity of OAS1 for the 5’-UTR of the viral genome, and in particular for the SL1 domain has been reported^1^’^2^. While the direct inhibition of OAS1 with small ligands can be difficult, this enzyme may be a good target for RNA therapeutics.^26^. Yet, despite these important perspectives the structural and molecular bases driving this selectivity remain elusive and poorly characterized. In this contribution by using equilibrium and enhanced sampling MD simulations, we have provided a rationalization of the SL1 tertiary structure and of its specific interactions with the OAS1 RNA recognition region. In particular we have evidenced that the particular features of the SL1 moiety, and in particular its high bending, induces a slightly different interaction network compared to ds-RNA. More importantly, in addition to the contact with minor groove areas the interaction between the protein and the nucleic acid is also driven by the interaction with the dangling extrahelical bases. This is particularly true concerning the loop area, that exhibits some flexibility and is also strongly interacting with polar protein residues; the peculiar role of H248 in strongly anchoring the nucleic acid fragment has been evidenced. As a matter of fact, the interaction involving the hydrophobic nucleobase will certainly provide a higher degree of selectivity than salt bridges mainly involving the backbone phosphates, hence providing a rationale of the observed preference of OAS1 for the SL1 region. Although, the full study of the interaction between OAS1 and 5’-UTR regions of other coronaviruses such as SARS-CoV or MERS,is out of the scope of the present contribution, the high sequence similarity, and the conservation of a similar SL, may suggest that our results can be extended to other viral strains. In the future we also plan to analyze the allosteric modulation of the OAS1 structure induced by the interaction with SL1. However, our results are important in providing an atomistic resolved vision on the reasons behind the recognition of SARS-CoV-2 genetic material by the innate immune systems, and hence can, in the long-term, help in designing therapeutic RNA strategy^26^ based on the stimulation of the immune system response by SL1 analogs.

The authors thank GENCI and Explor computing centers for computational resources. E.B. thanks the CNRS and French Ministry of Higher Education Research and Innovation (MESRI) for her postdoc fellowship under the GAVO program. AM thanks ANR and CGI for their financial support of this work through Labex SEAM ANR 11 LABX 086, ANR 11 IDEX 05 02. The support of the IdEx “Université Paris 2019” ANR-18-IDEX-0001 and of the Platform P3MB is gratefully acknowledged.

## Conflicts of interest

There are no conflicts to declare.

## Supplementary Materials for

### Computational details

The starting structure for SL1 was generated using the RNAcomposer webserver,^1^ and the OAS1 initial geometry was extracted from the crystal structure of the OAS1/ds-RNA complex (PDB ID 4IG8). The starting models for the OAS1/SL1 complex were generated by docking of representative structures from micro-second unbiased MD simulations of isolated SL1 and OAS1 using the HDock webserver.^2^ The model chosen as starting structure for the MD simulations was the best ranked by the HDock algorithm. The control system was taken as the above-mentioned crystal structure showcasing OAS1 in interaction with a 18-bp RNA duplex. Protein residues were modeled using the ff14SB amber force field,^3^ and bsc0+OL3 corrections were applied for RNA. The system was soaked in a cubic TIP3P water box with a 10Å buffer and potassium counter-ions were added to ensure a neutral total charge, resulting in systems from ~22,000 to ~60,000 atoms for the isolated SL1 and the complexes, respectively.

### Protein/RNA Docking

The protein/RNA Docking was performed using the HDock online webserver with standard parameters (http://hdock.phys.hust.edu.cn/), and using OAS1 crystal structure and the prominent conformation of SL1 obtained by our MD simulations. HDock is based on a hybrid docking algorithm of template-based modeling and free docking, which provides ideal performances as extensively benchmarked elsewhere. ^2^ We also checked the HDock accuracy by performing a re-dock of the crystal structure of OAS1 with the RNA double strand (See Figure S4) obtaining a Root Mean Square Deviation (RMSD) of only 0.4 Å for the backbone atoms.

### Molecular Dynamics simulations

MD simulations were carried out using NAMD3.^4^ The Hydrogen Mass Repartitioning method (HMR)^5^ was used to allow a 4 fs time step for the integration of the equations of motion. To prepare the system, 10,000 minimization steps were performed imposing positional constraints on the backbone atoms. 12 ns equilibration at 300K followed, during which the constraints were progressively released. The temperature was kept constant using the Langevin thermostat with a 1.0 ps^-1^ collision frequency, electrostatic interactions were treated using the Particle Mesh Ewald (PME) protocol.^6^ After equilibration, the conformational ensemble was sampled along 1-μs production run and structures were dumped every 40 ps. The conformational sample was further explored using Gaussian Accelerated Molecular Dynamic (GAMD) simulations performed using the NAMD module. To this end an harmonic repulsive potential, following a Gaussian distribution, was added to the dihedral angles of the protein and the RNA. The potential was added considering an upper limit of the standard deviation of 6 kcal/mol every 10,000 steps. GAMD was run for a total time of 1.2 *μs.* The obtained biased sampling was reweighted to take into account the effect of the biasing potential, following the procedure described in ref,^7,8^ and considering a free energy surface defined by the projection of the trajectory on top of the two main Principal Component Analysis (PCA) vectors describing the RNA dynamic. The two main PCA vectors are also reported in Figure S5.

### Structural Analysis

The cpptraj module of AMBER18^9^ was used to calculate distances and to perform the clustering analysis based on the of RMSD of the complex. Plots were generated using using the ggplot2 package of R^10^ and representations of the molecules were rendered by VMD.^11^

## Supplementary figures

**Figure S1.**
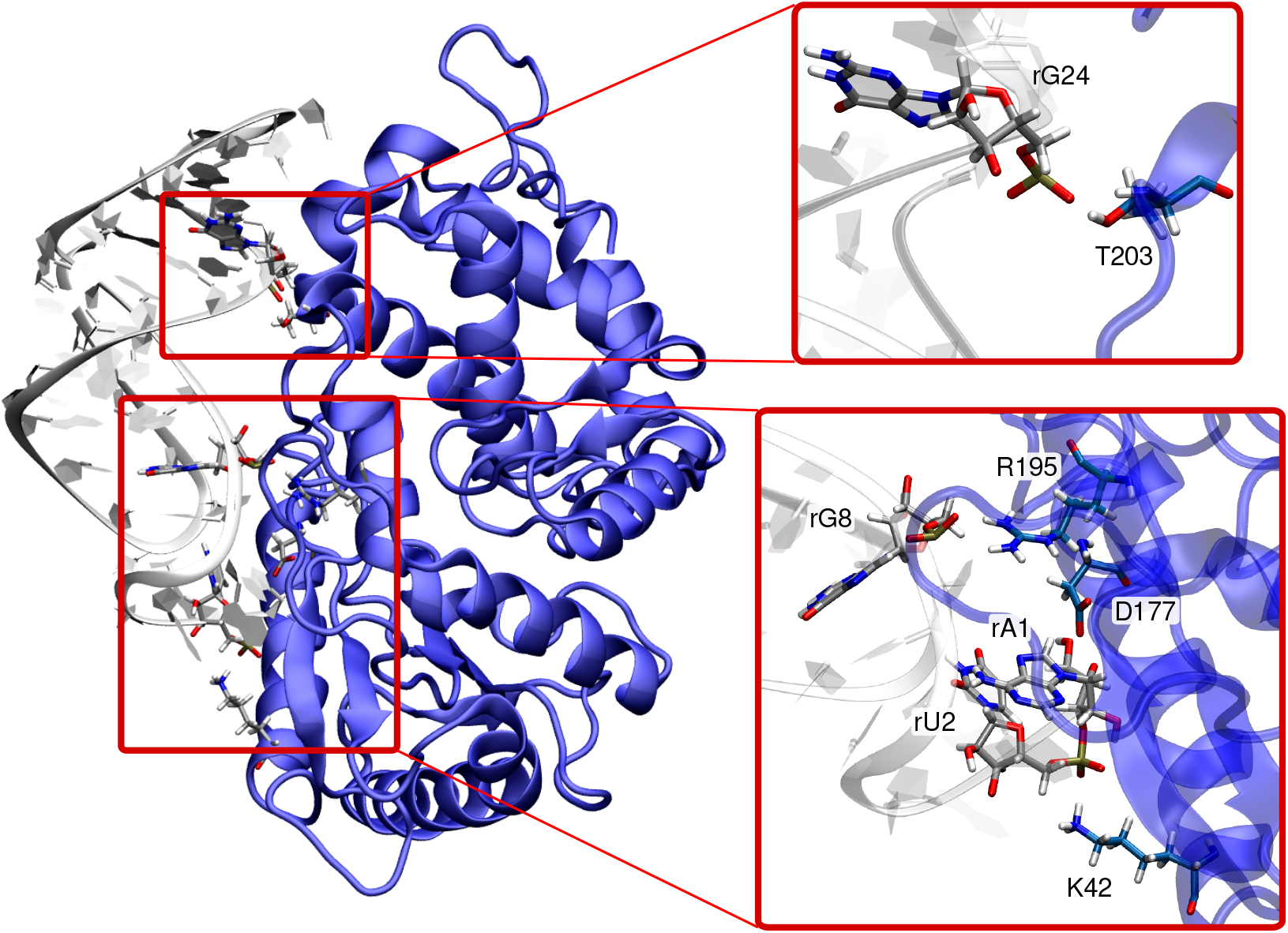
Representative structure of the OAS1/SL1 complex highlighting the contact surface with the RNA minor grooves. Magnified sections: zoom onto the key interactions at the two contact surfaces: T203:HG1-rG24:OP1, R195:HH12-rG8:OP1, D177:OD2-rA1:HO5’ K42:HZ1-rU2:OP2. Nucleic and amino acids carbon atoms are depicted in grey and blue, respectively.

**Figure S2.**
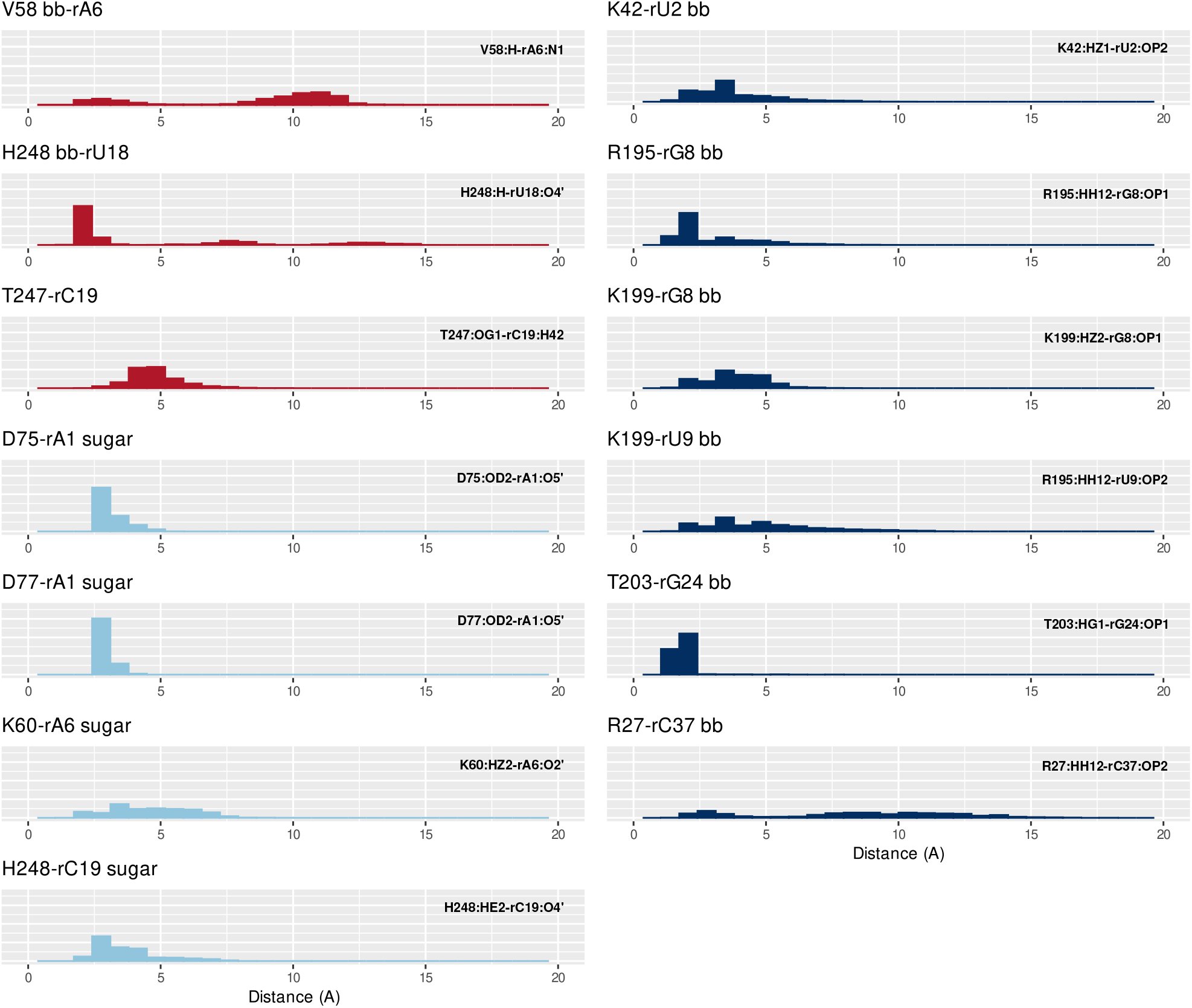
Distribution of the key-distances (in Å) involved in the OAS1/SL1 interaction network. Contacts with the nucleobases appear in red, with the sugar in cyan and with the backbone (bb) in blue. A label ‘bb’ is added behind OAS1 residue names when the interaction involves the backbone atoms and not the side chain.

**Figure S3.**
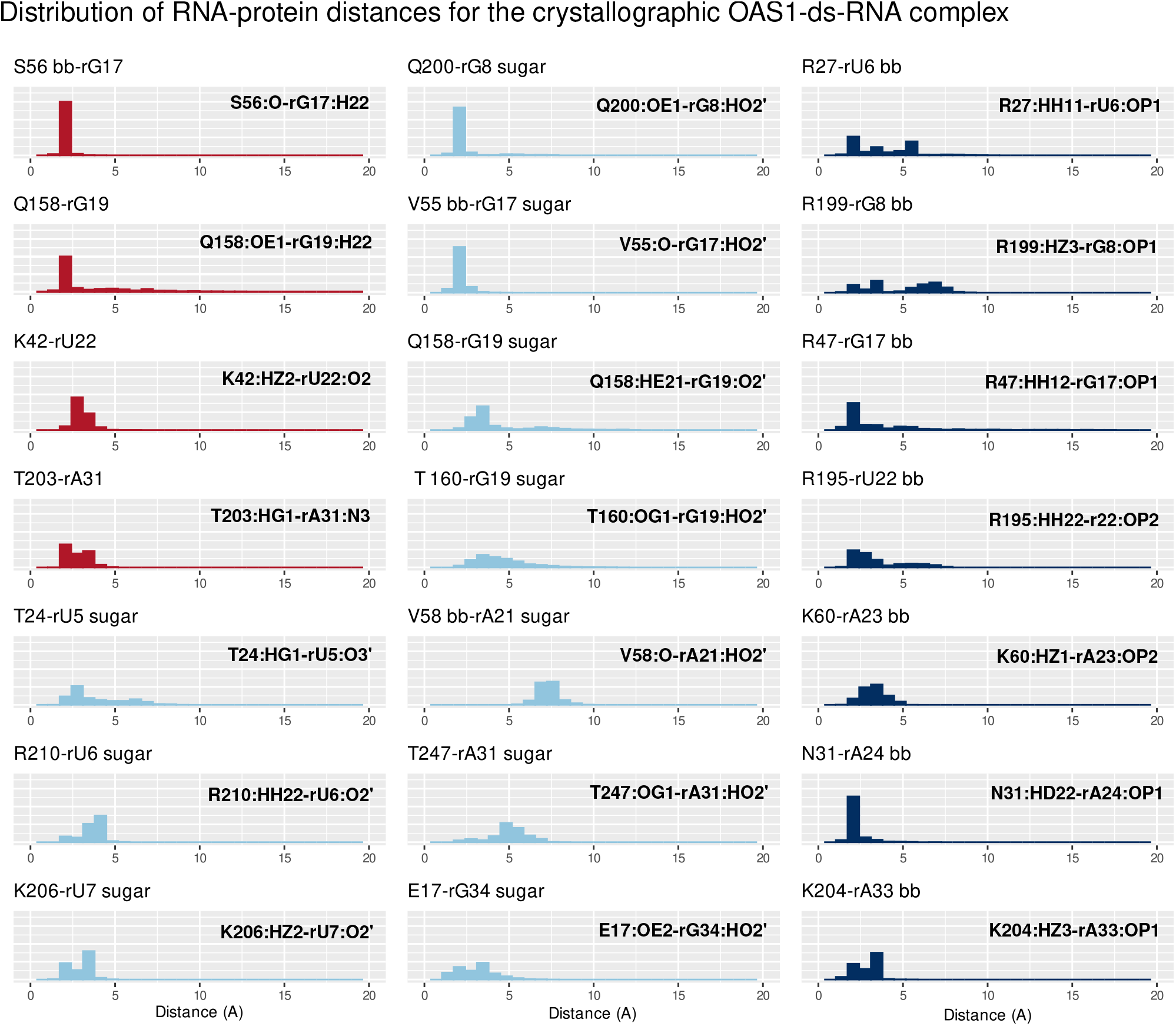
Distribution of the key-distances (in Å) involved in the reference system (OAS1/ds-RNA crystallographic complex) interaction network. Contacts with the nucleobases appear in red, with the sugar in cyan and with the backbone (bb) in blue. A label ‘bb’ is added behind OAS1 residue names when the interaction involves the backbone atoms and not the side chain.

**Figure S4.**
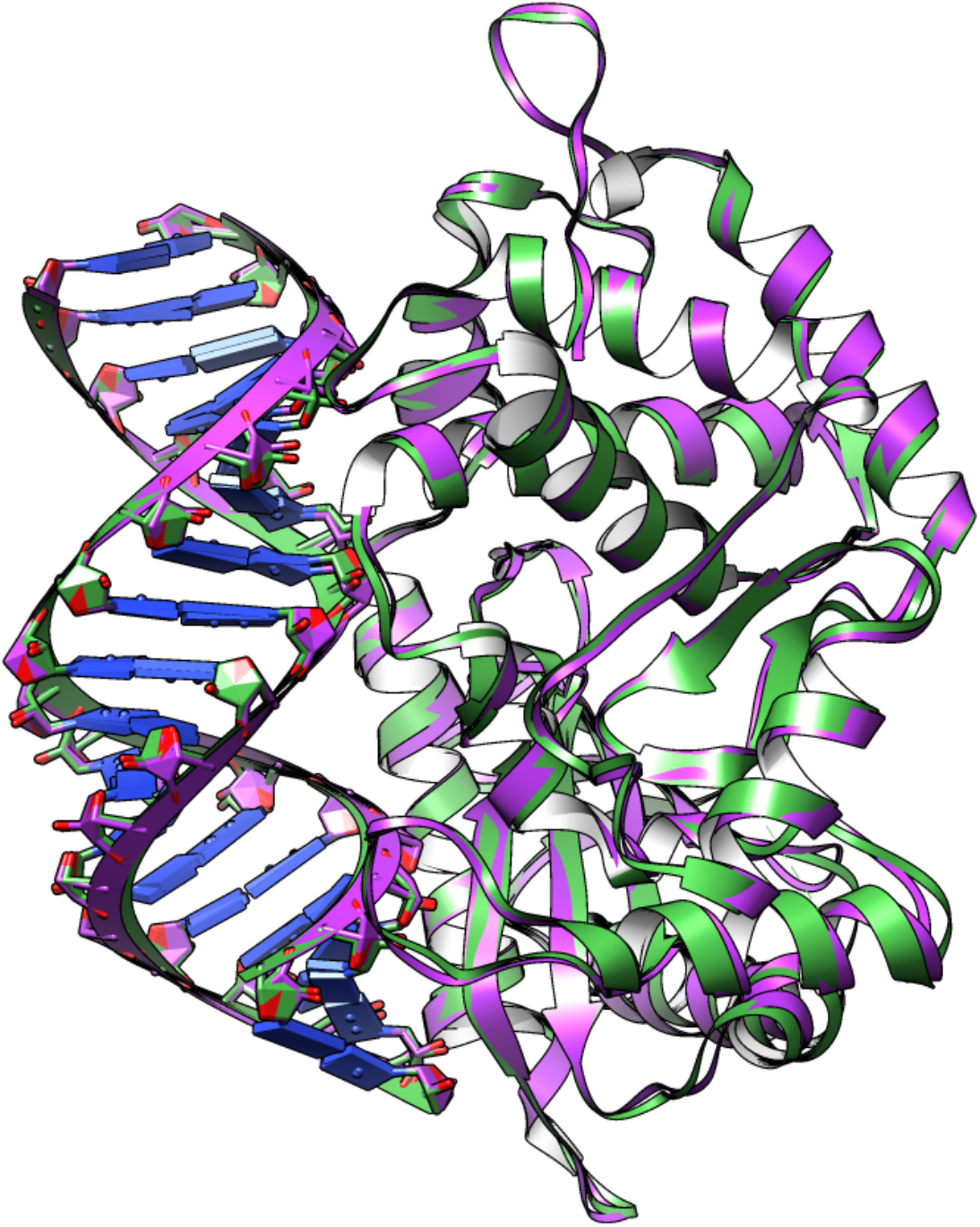
Superimposition of the dsRNA-OAS1 reference crystal structure (green) and the prediction of this complex by HDock (magenta), showing the very good performance of the latter.

**Figure S5.**
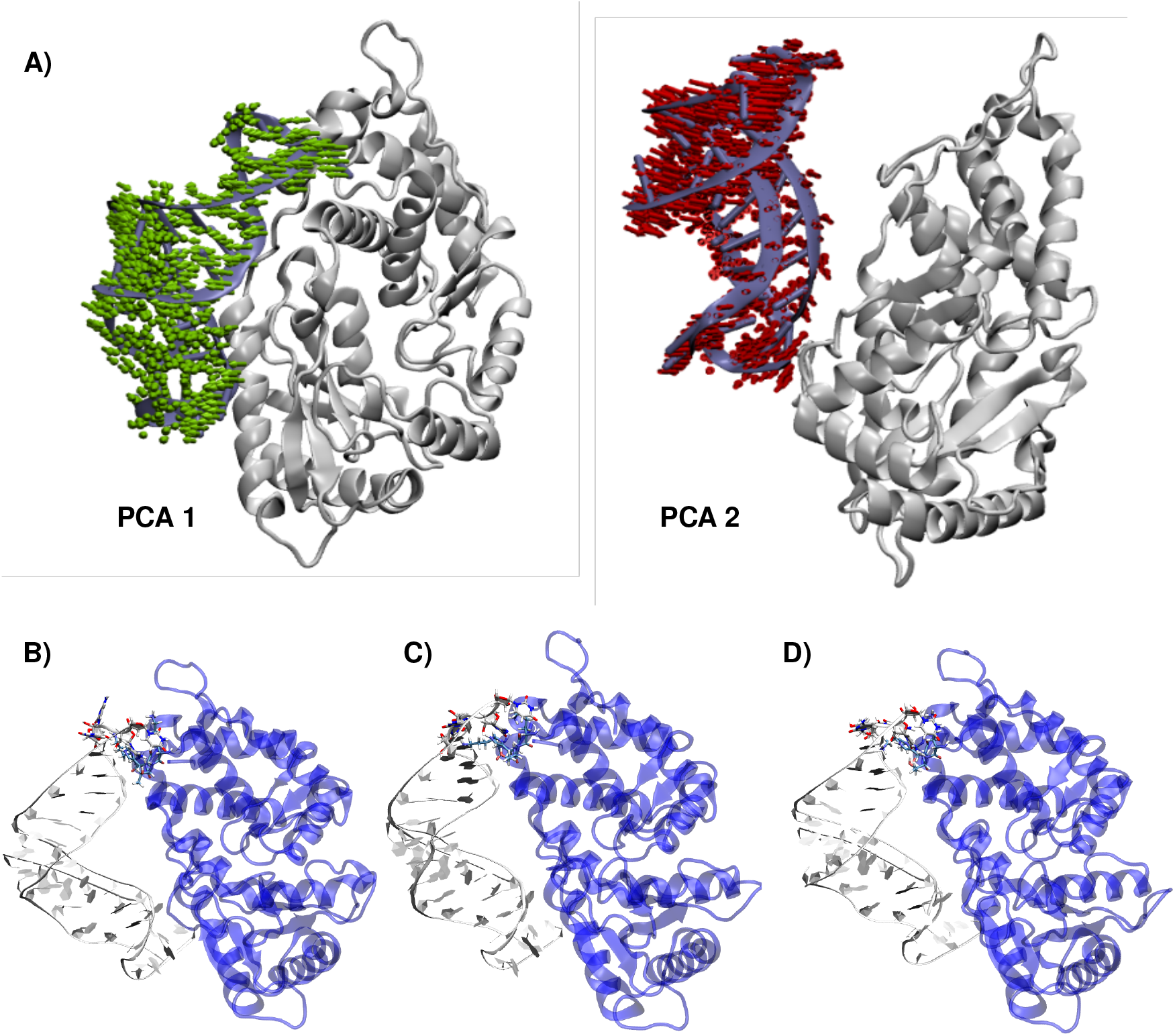
Top: Representation of the first (PCA 1) and second (PCA 2) collective modes issued by the PCA analysis and used as collective variables for the GaMD reweight procedure. Bottom: global view of the conformations corresponding to minimum energy regions labeled B), C) and D).

## Notes

### Competing Interest Statement

The authors have declared no competing interest.

